# A measure of vascular reactivity to overcome neurovascular uncoupling in functional imaging of brain tumors: initial results

**DOI:** 10.1101/240085

**Authors:** Henning U. Voss, Kyung K. Peck, Nicole M. Petrovich Brennan, Andrei I. Holodny

**Affiliations:** Department of Radiology, Weill Cornell Medicine; Department of Medical Physics, Memorial Sloan-Kettering Cancer Center; Functional MRI Laboratory, Department of Radiology, Memorial Sloan-Kettering Cancer Center; Brain Tumor Center, Memorial Sloan-Kettering Cancer Center

**Keywords:** BOLD fMRI, brain tumors, vascular reactivity, breath holding, neurosurgery

## Abstract

**Purpose:** Preoperative functional MRI (fMRI) is limited by a muted BOLD response caused by abnormal vasoreactivity and resultant neurovascular uncoupling adjacent to malignant brain tumors. We propose to overcome this limitation and more accurately identify eloquent areas adjacent to brain tumors by independently assessing vasoreactivity using breath-holding and incorporating these data into the BOLD analysis.

**Methods:** Local vasoreactivity using a breath-holding paradigm with the same timing as the functional motor and language tasks was determined in 16 patients (9 glioblastomas, 1 anaplastic astrocytoma, 5 low grade astrocytomas, and 1 metastasis). We derived a model based on coherence for analyzing BOLD fMRI that takes into account the altered hemodynamics adjacent to brain tumors.

**Results:** Activation maps computed using the coherence model were overall similar to standard activation maps. However, the coherence maps demonstrated clinically meaningful areas of activation that were not seen using the standard method in 12/16 cases. This included localization of language areas adjacent to brain tumors, where the coherence method results were confirmed by intra-operative direct cortical stimulation. Enhanced task response maps based on vasoreactivity mapping demonstrated more robust, anatomically-correct activation, in particular adjacent to tumors as compared to maps obtained without vasoreactivity information.

**Conclusions:** The present preliminary results demonstrate the principle that the neurovascular uncoupling known to affect the accuracy of BOLD fMRI adjacent to brain tumors may be, at least partially, overcome by incorporating an independent measurement of vasoreactivity into the BOLD analysis.

## INTRODUCTION

Neurosurgical resection remains the most important treatment option for malignant brain tumors as both the length and quality of survival are improved with maximized tumor resection [1,2]. Therefore, the goal of brain tumor surgery is to maximize the resection of the tumor while avoiding important adjacent eloquent cortices, whose inadvertent resection can lead to devastating neurological consequences. In order to preserve vital neurological function located near the tumor, it is important for the neurosurgeon to be able to identify the anatomical location of the eloquent cortices, either pre- or intraoperatively.

Traditionally, the eloquent cortices, such as the motor cortex, have been identified by direct cortical stimulation. More recently, blood-oxygen-level-dependent functional MRI (BOLD fMRI) has been used successfully in planning and carrying out the resection of brain tumors [3,4]. However, it has been found that BOLD maps adjacent to brain tumors have limited accuracy [5-9]. BOLD fMRI is based on the premise that there is a coupling between neuronal activity and blood flow. However, the neovasculature of malignant tumors is known to be abnormal both structurally and functionally [8], including changes in vascular reactivity [10-14]. Since BOLD fMRI measures the vascular response (rather than neuronal activation), in patients with malignant brain tumors the abnormal tumor neovascularity does not respond as vigorously to increased neuronal activity, which leads to a muted BOLD response [15]. A number of recent studies have correlated the anatomical location of neurovascular uncoupling, defined by quantitative measurements of cerebrovascular reactivity, with false negative BOLD fMRI results [7,16-18].

As a consequence, models assuming a uniform rather than an abnormal hemodynamic response function over the whole brain may not be sufficient to detect activation in an eloquent cortex influenced by the tumor. Data analysis based on the uniform hemodynamic response functions may lead to missed activation (or false negative errors) in the BOLD statistical parametric maps.

A possible way to overcome the limitation of false negative results caused by the presence of abnormal tumor neovasculature and resultant neurovascular decoupling is to independently measure vasoreactivity and to incorporate these data into the BOLD fMRI analysis. One way to independently measure vasoreactivity is by breath holding, which leads to hypoxia and hypercapnia, which under normal circumstances leads to vasodilatation of the cerebral vasculature [14,17,18]. These changes in vasodilatation can be measured by fMRI in the same way as BOLD fMRI responses to routine task paradigms are measured. The assumption is that the hemodynamic response function of abnormal neovasculature is altered from the norm in the same way irrespective of the stimulus (breath-holding or task fMRI). If this assumption holds true, then one should be able to detect BOLD activation even in areas of abnormal neovasculature by searching for coherence between the hemodynamic response function in the breath-holding and routine task fMRI paradigms.

The purpose of this study is to derive a voxel-specific model that takes into account altered hemodynamics by comparing BOLD response maps obtained during motor and language tasks with and without the incorporation of a breath-holding task. We hypothesize that BOLD response maps to motor and language tasks computed this way will be overall similar to standard activation maps based on uniform hemodynamic response models but will also show new areas of activation that accurately reflect increase in neuronal activity that were not detected by the standard method of BOLD fMRI analysis due to altered hemodynamics of the tumors.

## MATERIALS AND METHODS

The research performed was in full compliance with the Code of Ethics of the World Medical Association (Declaration of Helsinki). The institutional review boards of Memorial Sloan-Kettering Cancer Center and Weill Cornell Medicine approved the study. Informed Consent was not obtained, as the study was a retrospective review.

### Patient information

Each patient underwent an anatomical brain MRI and an fMRI as part of their routine preoperative care. All underwent resection of the tumor within two days of the MRI. Pathological examination of the resected tumors revealed the diagnosis of glioblastoma multiforme (*N* = 9), anaplastic astrocytoma (*N* = 1), low-grade glioma (*N* = 5), or metastasis (*N* = 1). See Table 1 for demographic data.

**Table 1:**
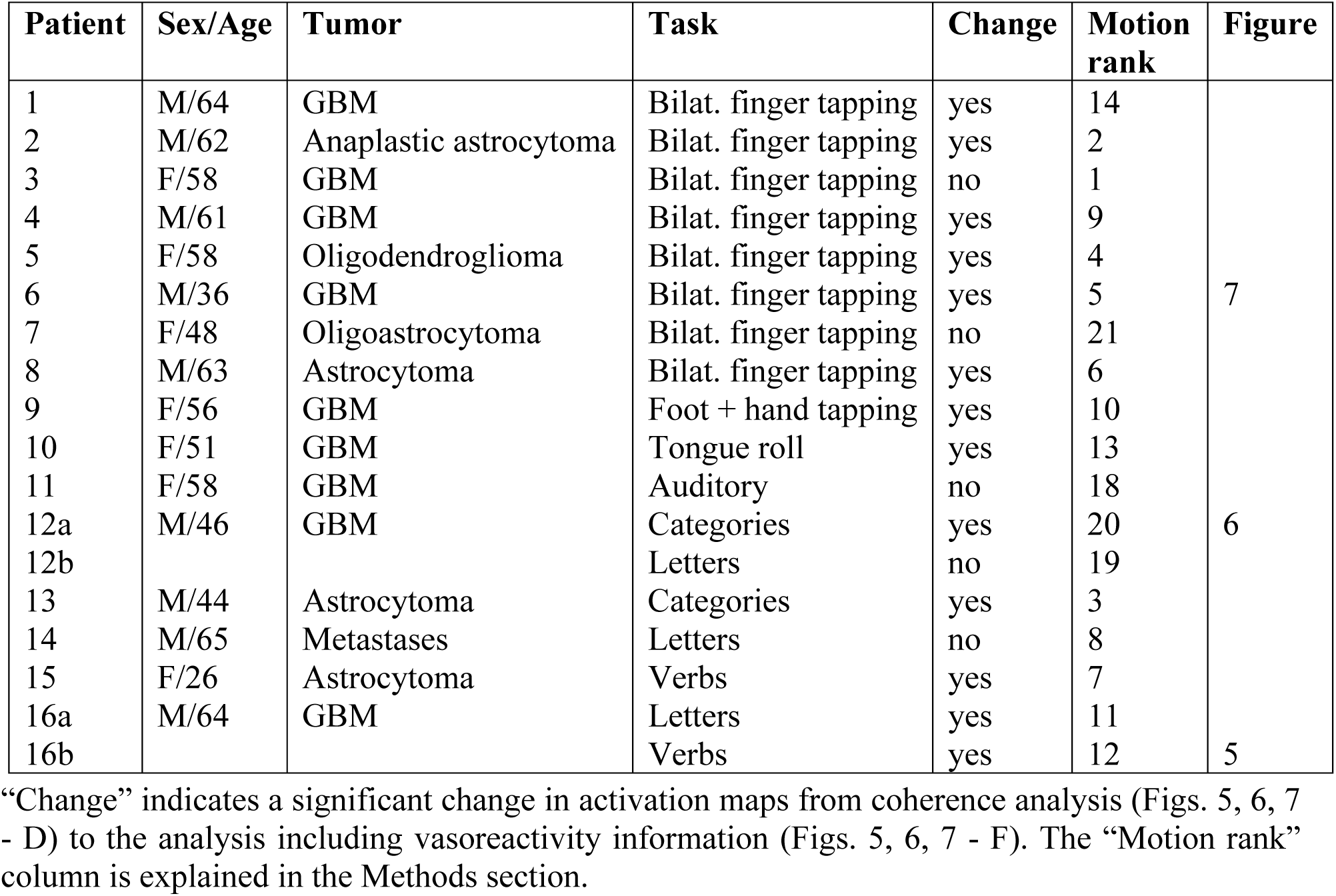
Patients, demographic information, etiology, and results.

### fMRI and anatomical MRI scanning

In total, 18 fMRI scans were performed on 16 patients. Patients also underwent an anatomical brain MRI on a 3.0 Tesla GEMS (Waukesha, WI) clinical MRI system with an 8-channel head coil. Breath-hold and task-specific fMRI were performed using echo-planar fMRI (TR = 4 s; TE = 40 ms; 90*^o^* flip angle; 128 × 128 matrix; 240 mm FOV; 4.5 mm slice thickness).

Six on/off blocks of 15 samples each were used in both breath-holding and task fMRI: 5 samples on (corresponding to inhalation and breath holding, or task, respectively), and 10 samples off (corresponding to normal breathing or resting/no task, respectively). During fMRI, patients either performed a motor and/or up to three language tasks, following aural instructions. See Table 1 for all tasks performed. Motor tasks consisted of bilateral finger tapping, foot and hand tapping, and tongue curling. In the finger tapping task the patient was asked to touch with the thumb consecutively the other four fingers in succession and start over again. It was self-paced at a rate of approximately two fingers per second, and had been practiced before the fMRI session. In the tongue curling task, patients were asked to make a small lateral movement of the tongue without opening the mouth. In the “foot and hand tapping” task, the finger tapping task was combined with foot-tapping. Language tasks consisted of “letters,” “verbs,” “auditory,” and “categories.” In the “letters” task, patients were asked to silently generate words that began with a given letter. In the “verbs” task, patients were presented with a noun and asked to generate action words associated with it. In the “auditory” task, patients were asked a variety of responsive naming tasks (e.g., what do you shave with?) and to answer the questions silently, avoiding overt verbalizations. In the “categories” task, patients were presented with a superordinate category and were told to generate subordinate words that fit the category. For breath holding, patients were instructed to take a deep breath (which took about 2 s) and then hold their breath for 18 s, before breathing normally again. Both breath-hold and task fMRI were acquired with 90 samples in total (6 on/off blocks of 15 samples), during 6 m of scan time, for each patient.

The severity of head motion in the breath-holding and the task scans was quantified by using the six rotational and translational parameter time series resulting from the three-dimensional co-registration procedure in AFNI. These motion parameters for the breath-holding and task scans were collapsed to a single number each in the following way: The mean squared sum of the three translational parameters was added to the mean squared sum of the rotational parameters for the breath holding scan, and then this sum was added to the sum of the equivalent task scan mean squared parameters. This was done for all 21 scans and the results were rank-ordered (small rank equivalent to relatively strong motion) and are provided in Table 1.

### fMRI data analysis

All analysis of fMRI and breath-hold echo planar imaging (EPI) data was done with AFNI [19] and MATLAB v. 2007b (The Mathworks).

Preprocessing was performed as follows: Motion correction, thresholding of brain versus background, voxelwise linear detrending of time series, and in-plane spatial smoothing with a Gaussian kernel of full-width-half-maximum = 3 voxels.

Subsequently, parametric maps were generated as follows: First, vascular reactivity maps were generated by correlating the breath-holding response time series *x_t_* voxelwise with the task indicator, or ideal, function *I_t_*, defined as *I_t_* = 1 for breath-holding or task samples and *I_t_* = 0 for normal breathing or rest samples. Secondly, parametric task maps were generated using three different methods: 1. Standard analysis based on the correlation of the BOLD signal with the ideal function, 2. Modified standard analysis based on the coherence between the BOLD signal and ideal function, and 3. Coherence-based analysis including vasoreactivity assessment from breath-hold data.

### 1. Standard analysis based on correlation coefficients

In this standard method, an ideal or task indicator function *I_t_* consisting of the numbers 1 and 0 was used to describe the timing of the motor and language paradigms. It was also used as a rough approximation of the hemodynamic response function template *h_t_*. The task activation maps were defined by correlating the signal in each voxel, *y_t_*, with the uniform ideal function, yielding a Pearson's linear correlation coefficient *R*(*y_t_, I_t_*). A significant positive task response was defined for *R*(*y_t_, I_t_*) > 0.55. This value corresponds to an average p-value of *p* = 1 × 10^−5^, set as an ad-hoc threshold in numerical simulations in order to compare with the coherence (see below). The one-sided tail probability for Pearson's linear correlation of *R*(*y_t_*, *I_t_*) = 0.55 is, however, *p* = 1.3 × 10^−8^. In other words, the latter p-value has been used as test statistics for determining single voxel significance. This is a quite conservative threshold even if the multiple testing problem is taken into account. In addition, a simple cluster-wise threshold was used; only clusters with at least two adjacent voxels of activation were retained in the final maps of correlation coefficients.

### 2. Modified standard analysis based on coherence

The spectral coherence between two signals is smaller than or equal to 1 and equals 1 for linear dependence. The maximum value of the coherence between *y_t_* and *I_t_*, *C*(*y_t_, I_t_*), in an interval [0.006 Hz, 0.028 Hz], centered at the block design task frequency of 0.017 Hz, was used as test statistics. The significance threshold corresponding to an average p-value of *p* = 1 × 10^−5^ was determined by Monte-Carlo simulations, and found to be *C*(*y_t_, I_t_*) = 0.79. Only clusters with at least two adjacent voxels of activation were retained in the final coherence maps.

### 3. Coherence based analysis incorporating vascular reactivity assessment from the breath-hold data

The conventional assumption in the most basic linear modeling of BOLD responses is the model

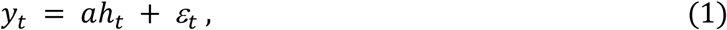

in which *h_t_* is the hemodynamic response, *a* is a coefficient to be estimated from *y_t_*, and *ε_t_* is noise not captured by the model. The hemodynamic response usually is considered to be independent of location, or uniform, throughout the brain. To take into account the possible dependence of *h_t_* on location in the brain, which is necessary to assess the impact of altered vascular reserve by the presence of tumors, here we assumed that *h_t_* varies depending on brain location and can be modeled from measuring the vasoreactivity *x_t_*, obtained from breath-holding scans with the same timing as the task in an additional fMRI experiment on the same patient. This assumption is motivated by the observation that the task response *y_t_* and the breath-hold response *x_t_* were very often simultaneously correlated or anti-correlated to *I_t_* in voxels in which significant task activation was observed (see Results). For this unknown dependence between task response and breath-hold response (vasoreactivity), we assumed the linear model

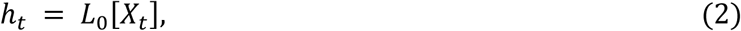

where *X_t_* is a local vasoreactivity template derived from the breath-hold signal *x_t_* as defined below and *L_0_* a linear functional. This relationship takes into account, for example, possible latencies between *x_t_* and *y_t_*. The task response model can now be written as

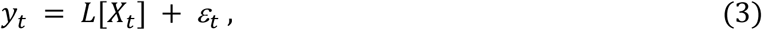

with again an unknown general linear dependence *L*. The degree of dependency can thus be estimated by the coherence between *X_t_* and *y_t_* [20], analogous to the second method. The regional vascular response template *x_t_* was computed by averaging the breath-hold signal *X_t_* over the six repetitions per experiment, for each voxel independently. The resulting signal of length corresponding to one breath-hold repetition was then concatenated six times to obtain a template time series *X_t_* of same length as the BOLD observations *y_t_* during the task experiment. Note that it is of crucial importance that the breath-hold and task block designs should match in length; here both paradigms consisted of 20 s “on” (corresponding to inhale/breath-hold and motor task, respectively) and 40 s “off’ (corresponding to normal breathing and rest), respectively. Analogously to the coherence based standard analysis, to generate activation maps, now the maximum value of the spectral coherence *C*(*y_t_, X_t_*) within the same frequency range as above was used. The same statistical thresholds for coherence as determined in the second step were used. Only clusters with at least two adjacent voxels of activation were retained in the final maps of coherencies. A pictorial representation of this method is seen in Fig. 1.

**Fig. 1:**
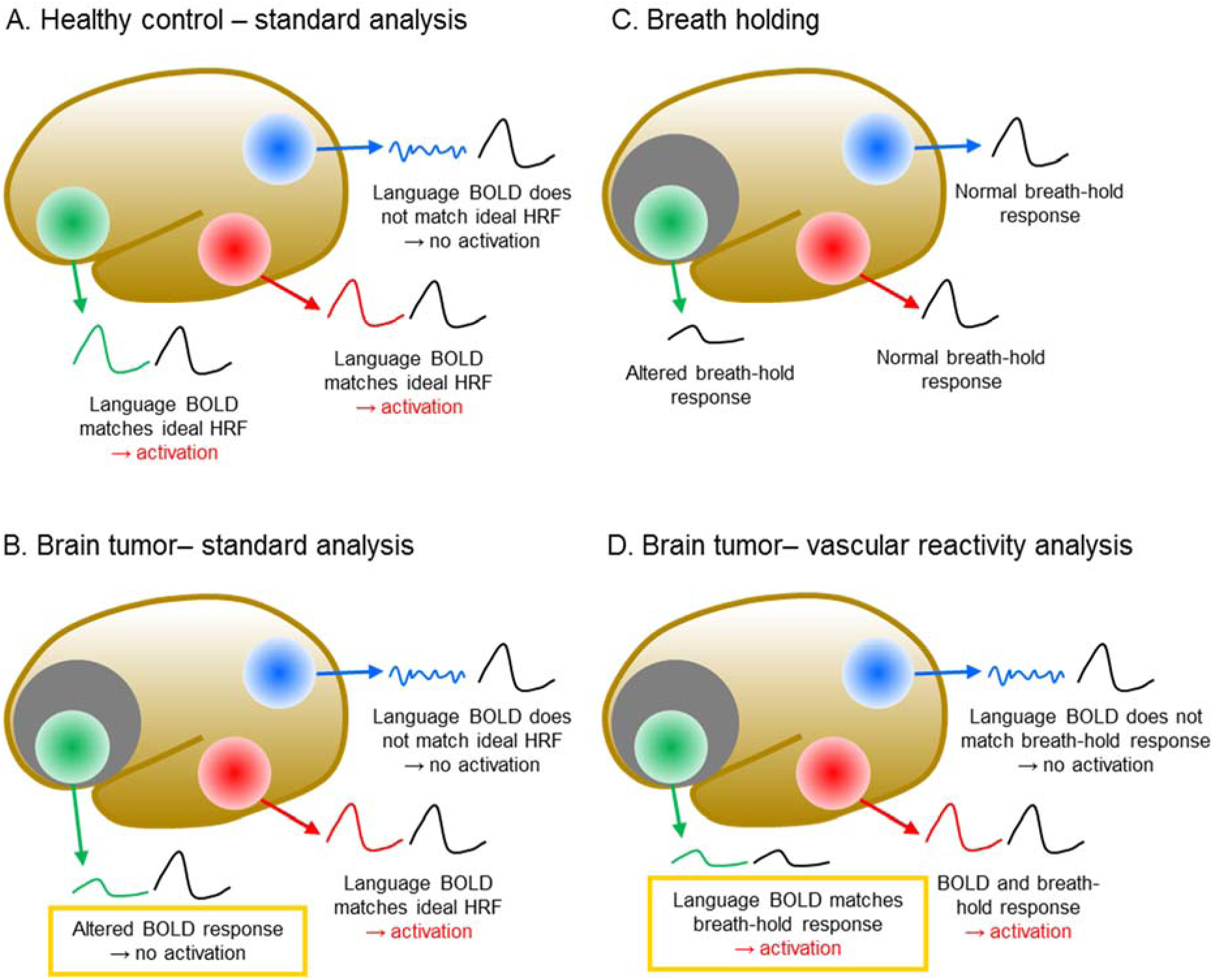
A schematic demonstrating the method used to incorporate breath holding data into the BOLD fMRI analysis. “A” represents the standard method of analysis of a language fMRI data set, where the BOLD fMRI data is compared to a standard ideal hemodynamic response function (HRF). The fMRI data in Broca’s area (Green) and Wernicke’s area (red) matches the standard HRF (in black), but the fMRI data in the occipital lobe (not related to language function) does not match the standard HFR. This method results in “activation” in Broca’s area and Wernicke’s area but not in the occipital lobe. In “B”, the shaded area represents a tumor with abnormal neurovascular reactivity (neurovascular uncoupling), which leads to a muting of the BOLD HRF. Consequently, the HRF in Broca’s area no longer matches the ideal HRF and Broca’s area is no longer “activated”. “C” demonstrates the HRF in each region during breath holding. In Wernicke’s area and in the occipital lobe, the breath holding HRF remains normal, but in Broca’s area, the breath holding HRF is muted. In the enhanced coherence method described in the present paper, instead of comparing the HRF obtained during the fMRI examination to an ideal, we compared it to the breath holding HRF-an independently obtained measure of the HRF. In the illustration, both Broca’s are and Wernicke’s area now demonstrate “activation”.

## RESULTS

### Relationship between the task response and the breath-hold response

We found that very often a significant positive task response was associated with either a positive or negative breath-hold response. This is demonstrated in Fig. 2, which shows scatterplots of typical correlation coefficients *R* of task response with task indicator function and breath-hold response with task indicator function for all voxels in a brain. Lines indicate significance thresholds. A significant task response was defined as |*R*| > 0.55 or an average p-value of *p* < 1 × 10^−5^, which is the same level as used for all analysis.

**Fig. 2:**
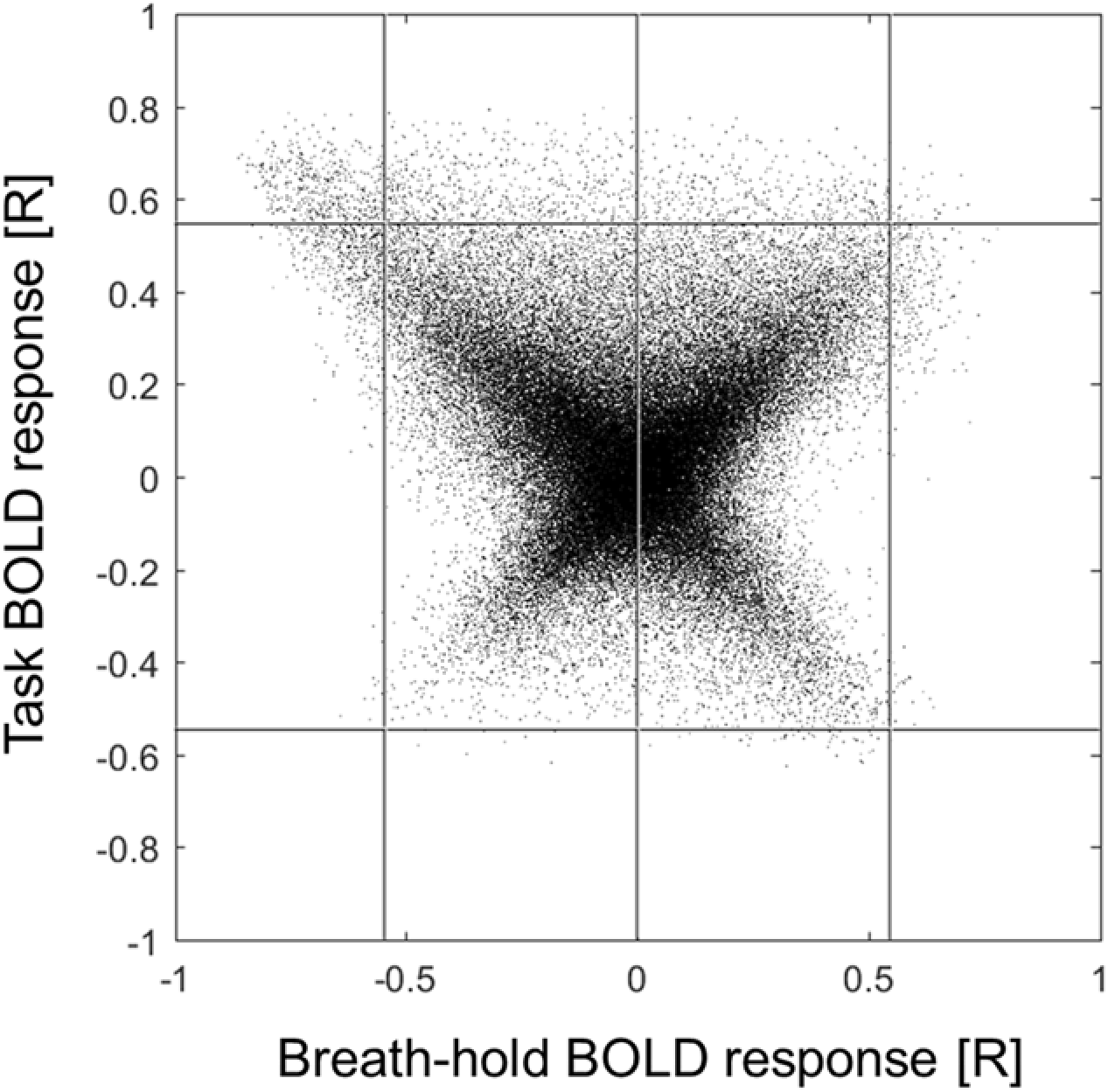
Relation between task and breath-hold response. Scatterplot of task and breath-hold BOLD response expressed as Pearson correlation coefficient *R* between the BOLD signal and the task indicator function for each voxel in the brain (Patient 7). The lines superimposed on the plot denote significance thresholds (*p* < 1 × 10^−5^) for the correlation coefficients (based on average values from Monte-Carlo simulations), both for positive and negative values each, and the zero line for breath-hold response to guide the eye.

Figure 3 shows scatterplots for all patients; in most cases a structure is visible, indicating a relationship between task and breath-hold responses.

**Fig. 3:**
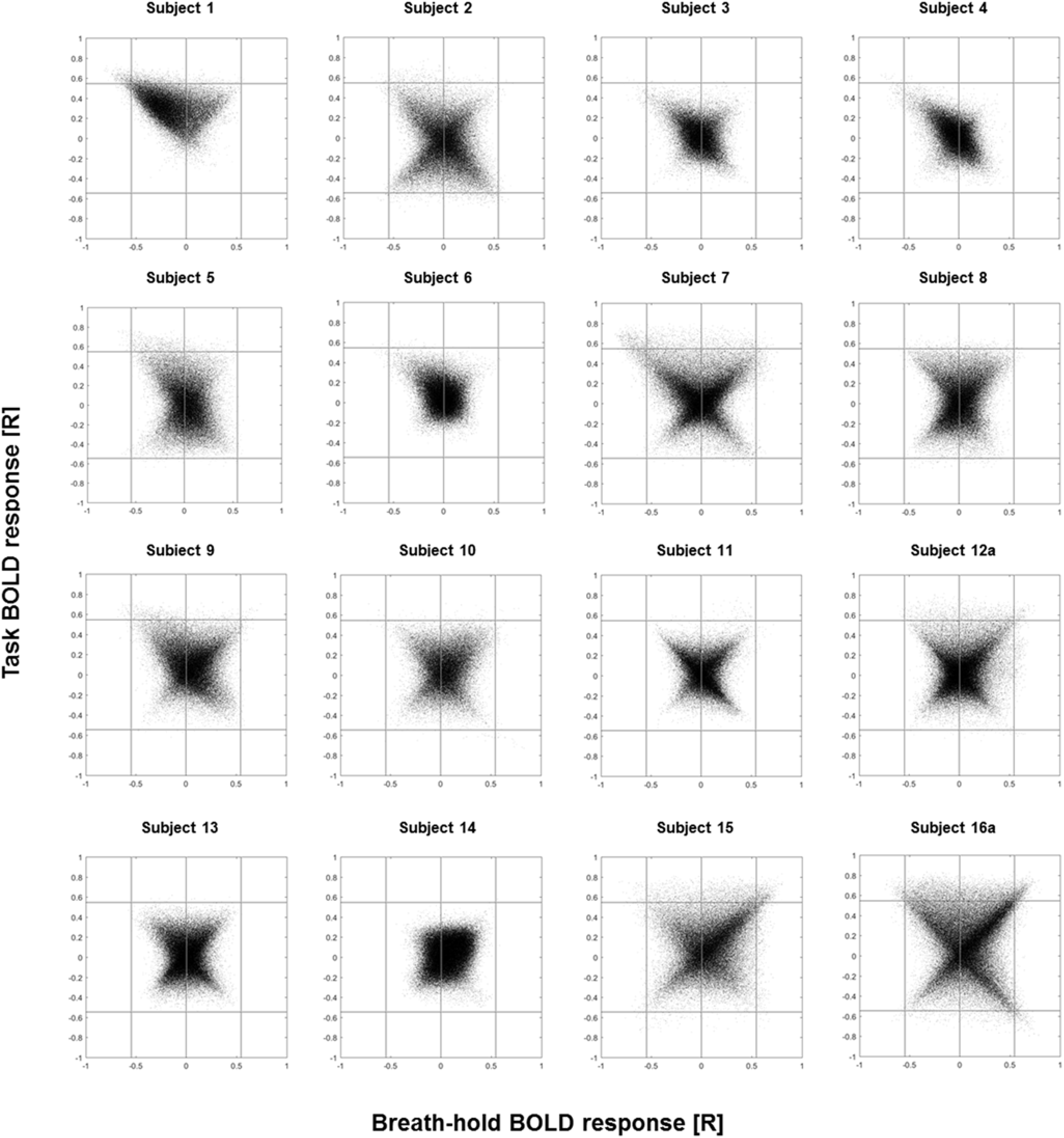
Relation between task and breath-hold response - all patients. Scatterplots of task and breath-hold BOLD response for all patients. Clear dependencies between task and breath-hold BOLD response correlation coefficients can be seen in all patients. For the sake of space, results for Patients 12b and 16b are not shown but look very similar to the shown first scan of the respective patient.

Figure 4A demonstrates the relationship between the task response and breath-holding template further; in the chosen voxel, breath-hold response and task response are strongly anti-correlated. The breath-hold response is also anti-correlated to the task indicator function, and the task response is correlated to the task indicator function; therefore, this voxel corresponds to one point in the upper left corner of Fig. 2. Because of the strong linear dependence between the task response and breath-hold template for this voxel, the coherence between both is expected to be similar to the coherence between the task response and task indicator function for the frequencies of interest. This is demonstrated in Fig. 4B. These observations motivated our approach to quantify task response using additional information from the breath-hold response.

**Fig. 4:**
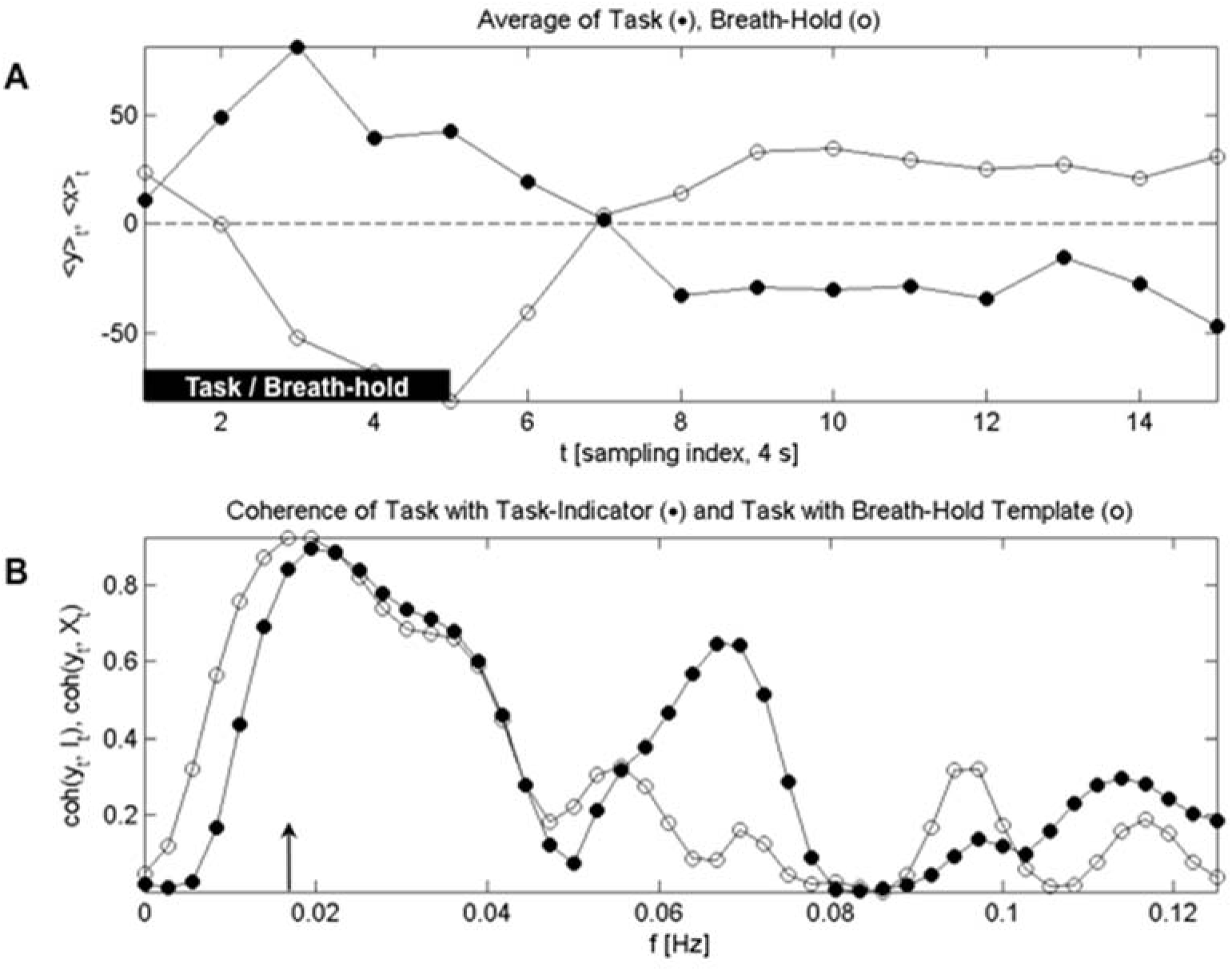
Task and breath-hold response. A: Example for task response (•) and breath-hold response (○). In this particular case, task response and breath-hold response are anti-correlated. The task and breath-hold period is marked by the black bar. B: This anticorrelation is also reflected in the high coherence at the frequency of the task and breath-hold paradigms (arrow). The maximum value of the coherence is slightly larger for the model based on local hemodynamic responses derived from breath-holding data (○) than for the standard model that uses a uniform hemodynamic response over the whole brain (•).

### Comparison of the standard method of analysis and the proposed coherence method

Review of the fMRI scans superimposed on the anatomical images demonstrated similar results between the standard method of analysis and the proposed coherence method. The only difference was that the coherence maps showed more areas of activation adjacent to the tumor. This was a retrospective study and in the actual clinical presentation of each specific case, the standard method was used; the coherence method was performed after the operation. Hence, we could only estimate the clinical utility of the proposed coherence method in a retrospective manner. Nevertheless, we judged that the new information made available from coherence analysis and particularly when vasoreactivity was incorporated (information which was not appreciated using the standard method) would have been clinically meaningful in 12/16 (75%) of the cases. This is illustrated by the following three examples:

### Example 1

Figure 5 demonstrates a case where activation near the tumor shows an atypical BOLD response that is picked up by the coherence method but not by the standard method. Incorporation of vasoreactivity map into the analysis caused a major change in activation. In the activation map using the standard correlation analysis, one can clearly see activation in Broca’s area ipsilateral to the tumor but the activation in the expected location of Wernicke’s area is bilaterally symmetrical and the relationship between Wernicke’s area and the tumor cannot be accurately established. Meanwhile, the coherence method clearly establishes the laterality of Wernicke’s area and its relationship to the tumor. The location of Wernicke’s area obtained in the post-operative period using coherence maps corresponded to the location of Wernicke’s area obtained by direct cortical stimulation.

**Fig. 5:**
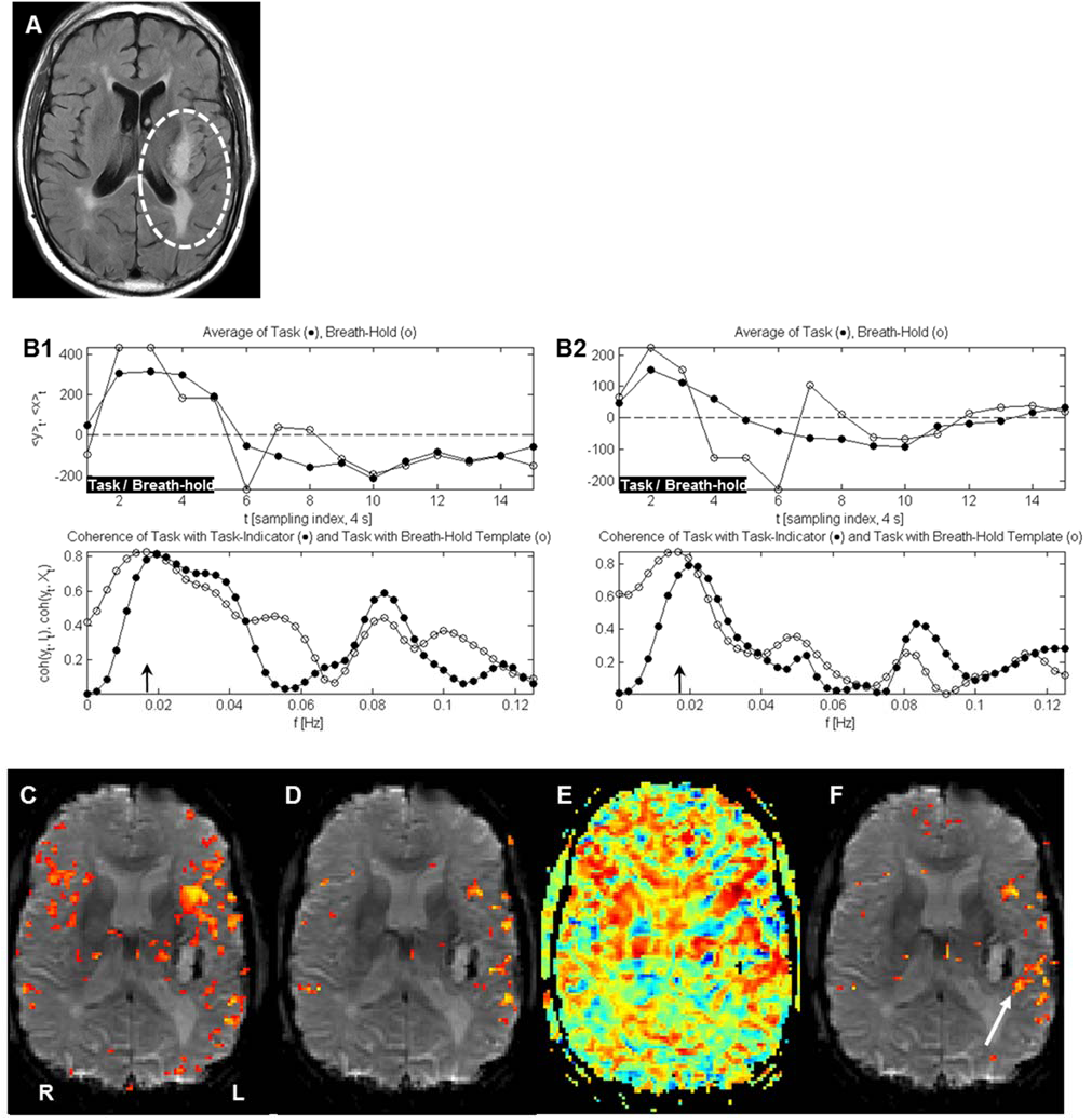
Example - Glioblastoma involving the left basal ganglia and temporal lobe. A: Anatomic image through the tumor (Patient 16b). B1: Breath-hold template and BOLD response to a language stimulus in a representative voxel in left Broca’s area, distant from the tumor (upper plot). Both responses are strongly correlated and cause similar coherence (lower plot) and activation maps (E, F). Therefore, at this position in the brain, the standard BOLD maps and the BOLD maps including cerebrovascular reactivity information look very similar. B2: Same for a voxel within Wernicke’s area, next to the tumor (arrow in Fig. F). Both BOLD response and cerebrovascular reactivity are altered as compared to the voxel remote from the tumor in Fig. B1. In addition, whereas the standard BOLD analysis gave a sub-threshold response, the coherence still demonstrates significant linear dependence between BOLD and the breath-hold template and thus is still able to indicate functional response. C: BOLD response (correlation map) in a language task (“verbs”). The correlation map fails to localize or lateralize Wernicke’s areas, i.e., there is no difference between the BOLD responses in the two hemispheres. D: The corresponding coherence map. E: Vasoreactivity map of the breath-hold response (blue negative R, red positive R, not thresholded). F: The activation map that utilizes information from vasoreactivity (coherence of the task response with the breath-hold template). Activation in Wernicke’s area is evident (arrow), which was not visible using standard analysis (C). Using the proposed calibration, Wernicke’s area was clearly identified and lateralized, which was a major preoperative concern to the neurosurgeon. Note the strong positive vasoreactivity (E) seen in this area.

### Example 2

Similarly, Fig. 6 shows a representative slice through the tumor of a patient with glioblastoma multiforme (GBM) (Fig. A), the relationship between vasoreactivity and task response for a language task for a specific voxel as indicated by the yellow arrow in Fig. F (Fig. B), and the BOLD response correlation map (Fig. C). The tumor is located anterior/superior to the dashed circle. Also see the corresponding coherence map shown in Fig. D. Figure E shows the vasoreactivity map of the breath-hold response (blue negative R, red positive R, not thresholded). Finally, Fig. F shows the coherence of the task response with the breath-hold template or, in other words, the map that utilizes information from vasoreactivity. Here, activation in Heschl’s gyrus (green and yellow arrows) and additional activation in Wernicke’s area (white arrow) are evident, which were both not visible in standard analysis (C) and weaker in the corresponding coherence map without breath-hold information (D). Note the strong positive vasoreactivity seen around areas of task-induced activation, and that both task and breath-hold responses look atypical since they are sustained only for a short time, shorter than a typical BOLD response (compare Fig. 4). Again, from the neurosurgeon’s perspective, the routine standard method failed to provide useful information—as language function was not lateralized and the relationship of the tumor to Wernicke’s area was not defined. On the other hand, the same data analyzed using the coherence method answered both clinical questions. The location of Wernicke’s area obtained in the post-operative period using coherence maps corresponded to the location of Wernicke’s area obtained by direct cortical stimulation.

**Fig. 6:**
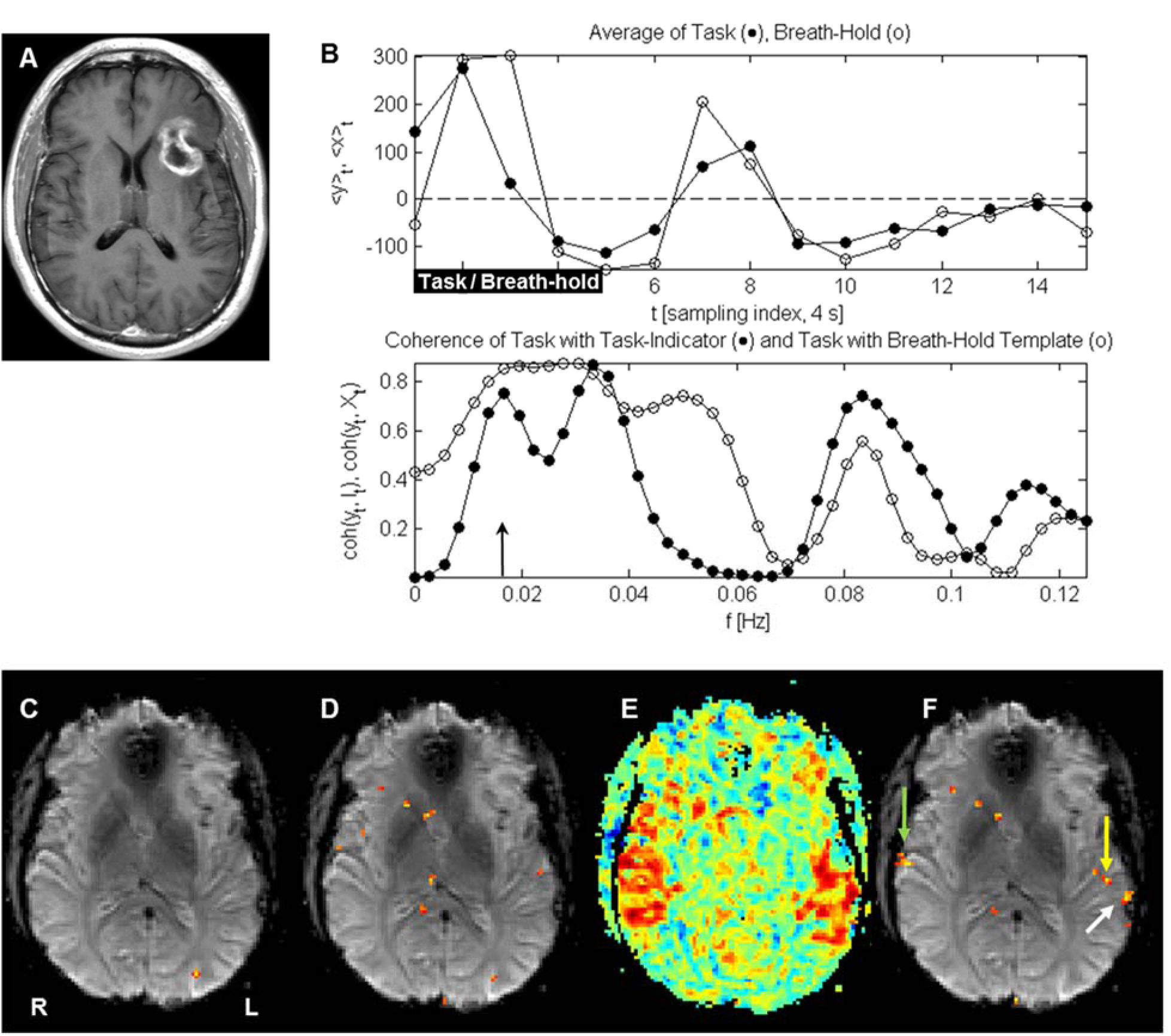
Example - Glioblastoma involving the left frontal and temporal lobe. A: Anatomic image including the tumor (Patient 12). The tumor is located superior to the EPI images in C to F. B, upper plot: Breath-hold template and BOLD response to a language task in one representative voxel in which the activation map obtained from vascular reactivity information shows additional activity, as indicated by the white arrow in F. In this case, task and breath-hold BOLD response are very similar, causing a high coherence between them in the frequency band centered around the task frequency (lower plot). Note that both responses look atypical since they are sustained only for a short time; a typical BOLD response would last longer (compare Fig. 4). C: BOLD response correlation map in a language task (“categories”). The correlation map failed to localize or lateralize Wernicke’s areas, i.e., there is no difference between the BOLD responses in the two hemispheres. Such a result is not helpful to the neurosurgeon planning the operation. D: The corresponding coherence map. E: Vasoreactivity map of the breath-hold response (blue negative R, red positive R, not thresholded). F: The activation map that utilizes information from vasoreactivity. Wernicke’s area is clearly identified and seen to be ipsilateral to the tumor (yellow arrow). In addition, bilateral auditory cortices (Heschl’s gyrus) are also clearly visualized (green arrow on the left). With the proposed calibration, the relationship of the tumor to Wernicke’s area becomes clear. This information is of crucial importance to the operating neurosurgeon. Identification of Wernicke’s area was not achieved using the standard method (C). This is an example in which positive vasoreactivity is observed, along with an atypical BOLD response shape. The additional BOLD response in the corrected map using vasoreactivity information arises from the fact that the coherency between breath-hold template and task response extracts the similarity between both curves (Fig. B, bottom plot), whereas the coherency between task response and task indicator is smaller due to the uncharacteristic shape of the task response.

### Example 3

The proposed coherence method was able to improve detection of eloquent cortices even at some distance from MRI defined enhancing tumor. Figure 7 provides another example for additional activation seen only if vasoreactivity information is utilized, again in a patient with GBM but in a bilateral finger tapping motor task. Whereas in the standard method motor activation did not exceed statistical thresholds in our high-resolution fMRI, the coherence method increased activations, and the further inclusion of vasoreactivity caused expected activation in the primary motor strip in both the ipsilateral as well as the contralateral hemisphere. Note the uncharacteristically long hemodynamic delay in the task response, which also shows up in the breath-hold response, and thus is detected by the coherence between them.

**Fig. 7:**
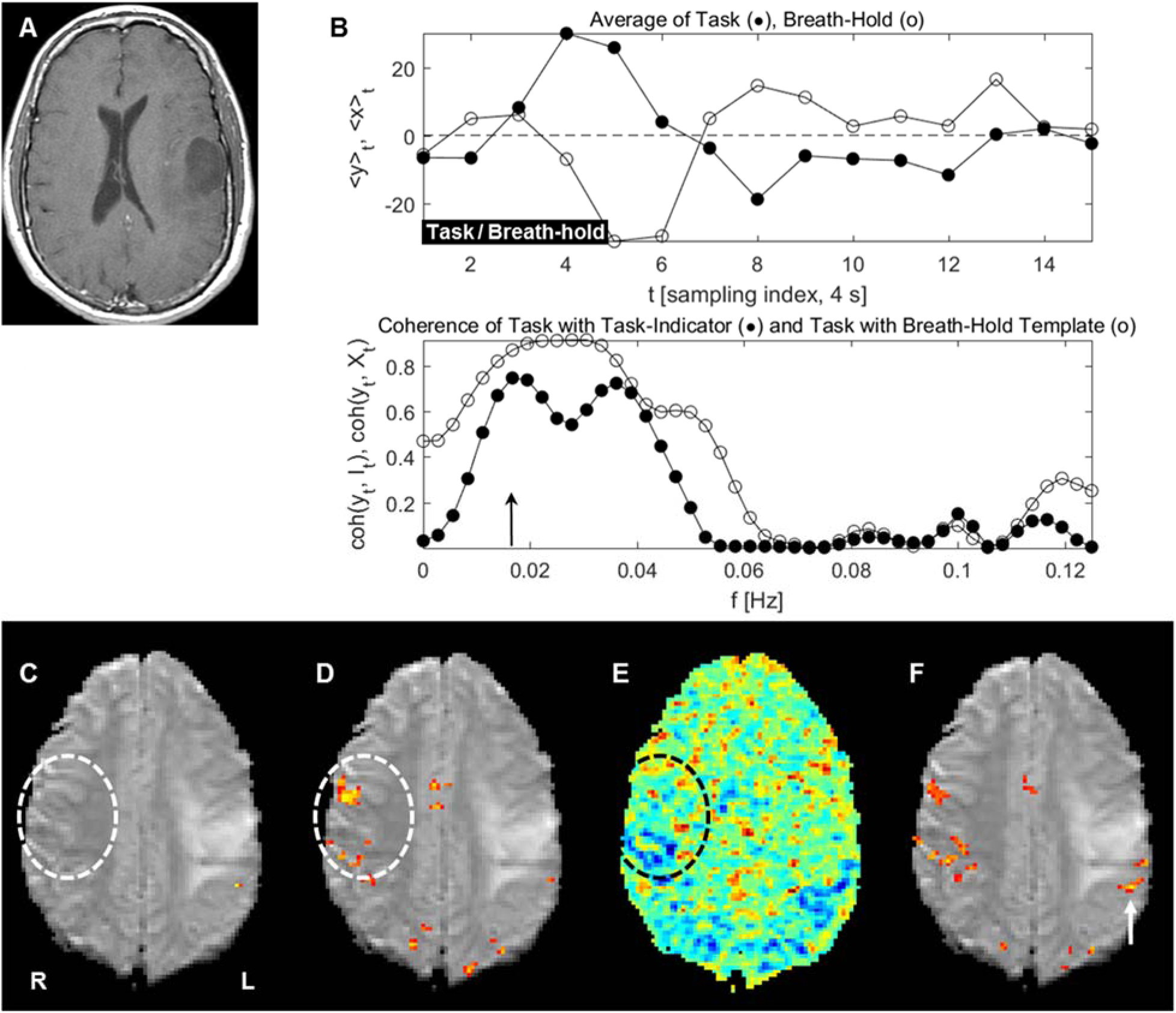
Example - Glioblastoma involving motor areas. A: Anatomic image with the tumor (Patient 6). B: upper plot: Breath-hold template and BOLD response to a motor task in a representative voxel in which the activation map obtained from vasoreactivity information shows additional activity, as indicated with an arrow in panel F. The lower plot shows the coherence between the task response and the breath-hold response with the task indicator function each. C: BOLD response in a bilateral finger tapping motor task using standard analysis. There is no activation in the motor area contralateral to the tumor and little activation adjacent to the tumor. D: The corresponding coherence map. E: Vasoreactivity map of the breath-hold response (blue negative R, red positive R, not thresholded). F: The activation map that utilizes information from vasoreactivity. Whereas in standard analysis motor activation did not exceed statistical thresholds in our high-resolution fMRI (C), coherence based analysis increases activations (D) and inclusion of vasoreactivity (E) causes expected activation in the primary motor strip (F) in both hemispheres. The inclusion of the vasoreactivity data, which entered our statistical analysis, enhanced the detection of motor activation in this area without causing any other spurious activation in the brain. The present example supports the contention that abnormal vasoreactivity and neurovascular uncoupling extends past the borders of the tumor defined on routine MRI sequences. The main reason for the detected response when vasoreactivity is utilized seems to be an uncharacteristically long hemodynamic delay in the task response, which also shows up in the breath-hold response, and thus is detected by the coherence between them, but not by the coherence between task indicator and task response, which has smaller values (B).

### Summary

In summary, we found significant changes in activation when the coherence method including vascular reactivity was performed. This finding may potentially lead to altering the neurosurgical decision-making process.

### Degree of motion in the breath-hold and the task scans

We also analyzed the degree of motion in the breath-hold and the task scans by using the six rotational and translational parameter time series resulting from the three-dimensional co-registration procedure in AFNI. These motion parameters for breath-hold and task scans were collapsed to a single number each, as described in the methods, to enable comparison between scans. The task motion was overall considerably smaller than the motion during breath holding; specifically, in 18 of the 21 scans, performed motion during breath holding was larger than during task.

## DISCUSSION

To map eloquent cortical areas in patients with brain tumors more precisely and to limit false negative BOLD fMRI results, we attempted to correct for altered hemodynamic response, which is characteristic of malignant tumors, by performing a simple breath-hold task and incorporating this information into the modeling of the BOLD fMRI response. The breath-hold BOLD response was often positively or negatively related to the motor or language task BOLD response in the same patient, depending on the region, as illustrated by scatterplots of correlation coefficients of both responses with the task indicator function (Figs. 2 and 3). A closer investigation of the BOLD response near a tumor in case examples showed various origins for altered hemodynamics, such as delays and shortened duration of responses.

The apparent ability of the proposed coherence method to identify eloquent cortices adjacent to brain tumors not seen with the standard method allows us to postulate the clinical utility of the proposed approach. We were only able to perform the coherence method retrospectively; however, our preliminary results in this study are convincing enough to pursue a future prospective study which would compare the standard method to our proposed coherence method and would use intra-operative cortical stimulation as the gold standard. For example, in Fig. 5, using the standard method, we were unable to define the language laterality to the operating neurosurgeon and we were unable to define the relationship of the tumor to the adjacent eloquent cortices (Wernicke’s area). However, when we retrospectively re-analyzed the case (after the operation had been performed) using the proposed coherence method, not only Wernicke’s area became clearly visible, but we were also able to unambiguously identify Heschl’s gyrus. Had this information been available to the operating neurosurgeon prior to the operation, it would have proved invaluable by clearly identifying the language lateralization (in this case, ipsilateral to the tumor) and the exact anatomical relationship between the tumor and the functional language area to be avoided during tumor resection. This information would have indicated the necessity for “awake mapping” during the operation, which would have required awakening the patient for anesthesia and asking the patient to perform various language paradigms during direct cortical stimulation to identify Wernicke’s area. The accurate localization of Wernicke’s areas by the proposed coherence method would have guided the direct cortical stimulation, probably decreasing the duration of the operation and possibly also decreasing the possibility of seizure onset caused by repeated cortical stimulation.

In the example in Fig. 5, the standard method was able to identify and lateralize Broca’s area (Fig. 5C). However, this information was of limited clinical utility as Broca’s area was at some distance from the tumor. The standard method showed symmetrical activation between the two hemispheres in the expected location of Wernicke’s area. Hence, we were unable to convey to the neurosurgeon any clinically useful information about 1) the relationship of the adjacent essential language area (Wernicke’s area) to the tumor, 2) the lateralization of Wernicke’s area, or 3) the possibility of cortical reorganization (plasticity) of language function in this patient due to the growth of an adjacent tumor [9]. Employing our proposed coherence method, we were able to answer all of the important, clinical, neurosurgical questions that were not answered with the standard method. Specifically, the laterality of Wernicke’s area is unambiguously established-ipsilateral to the tumor. The anatomical relationship of the tumor to Wernicke’s area was clearly displayed and the possibility of cortical reorganization was discounted.

In one case, we were able to identify activation in an eloquent cortex contralateral to the FLAIR-defined tumor (Fig. 7). Analysis of the bilateral finger tapping task using standard methodology failed to elicit activation in the expected location of the finger homunculus in the pre-central (motor) gyrus. However, analysis using the proposed coherence method was able to accomplish this task. Since pathologically defined malignant tumor cells and abnormalities in MRI breath-hold characteristics extend large distances past the borders of the MRI-defined gliomas [21], the case examples in this study raises the intriguing possibility of the correction of false negative BOLD fMRI results using the here proposed coherence method even at a distance from the MRI defined borders of the tumor.

In this study, we did not investigate whether altered hemodynamics correlated with proximity to the eloquent areas of the tumor, the pathology, or the related task. All analyses were performed retrospectively; therefore, we were not able to definitively correlate the results obtained to direct intra-operative cortical mapping of eloquent cortices. More experiments are necessary to investigate these concerns. However, overall, our results thus far suggest that the coherence method, especially with the inclusion of vasoreactivity data, may enhance functional mapping in patients with compromised hemodynamics secondary to pathology. To better evaluate our results, our future prospective study will include deriving heuristic methods to assess improvement in sensitivity or specificity.

The vasodilatory effects of varying CO_2_ content may be utilized in various ways to calibrate and improve functional BOLD detection [22,11,5,6,21,23]. Our proposed coherence method differed from previously used standard analysis, which were developed in healthy subjects, by utilizing our observation that breath-hold vasoreactivity and BOLD response are often linearly dependent. Our proposed coherence method did not put any effort into finding the exact relationship between them, which may be very difficult in vaculature altered by tumors. We think that due to the large number of possibilities of how the vascular response can be altered by a tumor, for example already by the location of the eloquent cortical areas with respect to the tumor, this much more ambitious goal would require a much larger study with many patients per tumor category, tumor location and extent, type of eloquent area, etc. Rather, our analysis was soelely based on the assumption that a linear relation between breath-hold vasoreactivity and BOLD response was a good or at least often useful approximation to the unknown true relationship. Of note, we found that both positive and negative vasoreactivity can occur in regions near or in tumors, where task response was also observed. We were neither aiming at separating response from the capillary bed from venous response for the tumor case. Our assumption of a linear relationship between vasoreactivity and response to a task was quite general but still amenable to modelling, for example via coherence analysis.

The effectiveness of the proposed coherence method relied heavily on the compliance of patients with both the breath-hold and the performed motor or language tasks. Breath-hold compliance can be very low in the clinical setting [4].Therefore, care had to be taken to ensure breath holding. In our study, we believe that all 16 patients indeed did perform a satisfactory breath-hold, based on the vasoreactivity data (Fig. 2). Another problem was head motion, often enhanced in tumor patients, by difficulties in performing motor tasks with keeping the rest of the body perfectly still, or language tasks without talking. Motion may affect both task response and vasoreactivity mapping, either causing false positive or false negative activations. In our study, motion was summarized as the relative rank of severity of motion for each patient in Table 1. We would not exclude the possibility that motion caused false activations in particular for patients with a low rank in Table 1.

In presurgical planning fMRI, as in any other fMRI study, it is imperative to use an adequate statistical threshold for the statistical parametric maps. Whereas in neuroimaging studies with normal subjects statistical thresholds are often set to a fixed value for all subjects encountered in a group study, for clinical patients with brain tumors an adequate choice of thresholds requires more thought. One reason is that often a considerably altered hemodynamic response weakens the BOLD signal observed, as has been demonstrated recently in pre-motor areas of clinical patients performing a motor task [7]. Another reason is that there is an on-average stronger head motion which reduces signal, since clinical studies cannot be performed in the same controlled way as research studies and also have to take disease-related task performance limitations such as fatigue into account. Whereas this does not mean that statistical thresholds can be set arbitrarily (in particular so this does not lead to too many false negatives or to miss BOLD responses), often in clinical practice thresholds are chosen less conservatively than in research studies. In order to provide an unbiased result, in the present study, thresholds were set to a constant significance value of *p* = 1 × 10^−5^; the corresponding threshold for the correlation coefficient is *R* = 0.55 and for the coherence *C* = 0.79 (as determined by numerical simulation). We chose this rather conservative approach to have a common basis for comparisons; it might be that more differences between standard maps and maps based on vasoreactivity become evident once the thresholds are set individually for each case.

We based our analysis on the well-known vasodilatory effects of CO_2_ in brain vasculature, which may be observed with BOLD fMRI imaging. We attempted the first steps towards the goal of understanding how vasoreactivity can be utilized towards a better definition of neuronal response in patients with brain tumors. We used the relatively crude and not well-controlled method of breath holding to measure vasoreactivity. Whereas breath holding is very easy to apply in the clinical population without the need of additional equipment and personnel, other more direct methods of hypercapnia, for example by delivering pre-mixed gas [24], may be better controlled and may lead to better results if combined with fMRI.

## CONCLUSION

Preoperative BOLD fMRI is limited by a muted BOLD response caused by abnormal vasoreactivity and resultant neurovascular uncoupling adjacent to malignant brain tumors. We proposed to overcome this limitation and more accurately identify eloquent areas adjacent to brain tumors by independently assessing vasoreactivity using breath holding and incorporating these data into the BOLD standard method of analysis using coherence. The generated coherence maps demonstrated clinically meaningful areas of activation that were not seen using the standard method of analysis in 12/16 cases. This included localization of language areas adjacent to brain tumors, where results using coherence analysis were confirmed by intra-operative direct cortical stimulation. The present preliminary results demonstrate the principle that the neurovascular uncoupling known to affect the accuracy of BOLD fMRI adjacent to brain tumors may be, at least partially, overcome by incorporating an independent measurement of vasoreactivity into the BOLD standard method of analysis. These initial, limited results need to be substantiated by further, prospective studies.

## Acknowledgments

This research is funded in part through the NIH/NCI Cancer Center Support Grant P30CA008748 and the NIH/NCI R21CA220144-01 award.

